# Development of the 12-Base Short Dimeric Myogenetic Oligodeoxynucleotide That Induces Myogenic Differentiation

**DOI:** 10.1101/2024.03.14.584904

**Authors:** Koji Umezawa, Rena Ikeda, Taiichi Sakamoto, Yuya Enomoto, Yuma Nihashi, Sayaka Shinji, Takeshi Shimosato, Hiroshi Kagami, Tomohide Takaya

## Abstract

A myogenetic oligodeoxynucleotide (myoDN), iSN04 (5’-AGA TTA GGG TGA GGG TGA-3’), is a single-stranded 18-base telomeric DNA that serves as an anti-nucleolin aptamer and induces myogenic differentiation, which is expected to be a nucleic acid drug for the prevention of disease-associated muscle wasting. To improve the drug efficacy and synthesis cost of myoDN, shortening the sequence while maintaining its structure-based function is a major challenge. Here, we report the novel 12-base non-telomeric myoDN, iMyo01 (5’-TTG GGT GGG GAA-3’), which has comparable myogenic activity to iSN04. iMyo01 as well as iSN04 promoted myotube formation of primary-cultured human myoblasts with upregulation of myogenic gene expression. Both iMyo01 and iSN04 interacted with nucleolin, but iMyo01 did not bind to berberine, the isoquinoline alkaloid that stabilizes iSN04. Nuclear magnetic resonance revealed that iMyo01 forms a G-quadruplex structure despite its short sequence. Native polyacrylamide gel electrophoresis and computational molecular dynamics simulation indicated that iMyo01 forms a homodimer to generate a G-quadruplex. These results provide new insights into the aptamer truncation technology that preserves aptamer conformation and bioactivity for the development of efficient nucleic acid drugs.

**Key Contribution:** This study reports the structure-based shortening of a myogenetic oligodeoxynucleotide, iSN04, as an anti-nucleolin aptamer that induces myogenesis. The shortening technology of aptamers while maintaining their conformation and activity improves their potency of drug function and synthesis cost.

## 1. Introduction

Myogenic progenitor cells, called myoblasts, differentiate into myocytes, fuse into multinucleated myotubes, and eventually form myofibers during skeletal muscle development and regeneration. Myogenic differentiation is thus an initial key step for muscle growth and maintenance [1]. However, aging, chronic kidney disease, cancer, or diabetes mellitus decrease the differentiation capacity of myoblasts [2-5], which is one of the reasons for age- and disease-associated muscle loss such as sarcopenia and cancer cachexia. Therefore, molecules that enhance myogenesis may be promising drugs to prevent the muscle wasting that is increasing in aging societies. Oligodeoxynucleotides (ODNs) have been studied as nucleic acid aptamers that bind to their targets in a conformation-dependent manner with high specificity and affinity, similar to antibodies. Nowadays, aptamers are one of the potential next-generation drugs [6]. We have recently reported a series of myogenetic ODNs (myoDNs), which are single-stranded 18-base telomeric phosphorothioated (PS)-ODNs, that robustly induce myogenic differentiation of myoblasts [7] and rhabdomyosarcoma [8]. One of the myoDNs, iSN04 (5’-AGA TTA GGG TGA GGG TGA-3’), serves as an aptamer that physically interacts with nucleolin, a multifunctional phosphoprotein localized in myoblast nuclei [7,9]. Antagonizing nucleolin with iSN04 reverses nucleolin-inhibited translation of p53 mRNA and improves p53 protein levels, resulting in enhanced myogenesis [7]. Since iSN04 restores myoblast differentiation deteriorated by diabetes mellitus [10] or cancer-secreted factors [11], it is expected to be a drug seed to reverse muscle loss in these diseases. In addition, nucleolin inhibition by iSN04 suppresses inflammatory responses [12] and induces myocardial differentiation of pluripotent stem cells [13] by modulating the β-catenin signaling pathway. These results suggest that iSN04 may be applicable to inflammatory muscle diseases such as dystrophy and to cardiac regeneration therapy.

iSN04 is incorporated into myoblasts without a carrier [7]. In general, single-stranded PS-ODNs can be spontaneously incorporated into the cytoplasm by an endocytic process termed gymnosis, in part due to their lower molecular weights compared to double-stranded nucleotides [14]. Since shorter PS-ODNs are more favorable for intracellular uptake, sequence shortening is important pharmaceutically to improve drug efficacy and also beneficial industrially to reduce synthesis costs [15]. In this study, we attempted to shorten iSN04 while maintaining its myogenetic activity. In the previous report, computational simulations and mutational experiments revealed that the guanine stack (G-stack; GGG at the 13-15th bases) within iSN04 is the core structure. Interestingly, an isoquinoline alkaloid, berberine, binds to the G-stack and enhances the activity of iSN04 [7,9]. By forming a complex, berberine is thought to shift the conformation of iSN04 to a more stable and active form, suggesting that the ODNs that configure the G-stack may serve as myoDNs. Here, we investigated the myogenetic activities of the 11∼16-base non-telomeric PS-ODNs predicted to form the G-stack structure. Several PS-ODNs successfully induced human, mouse, and chicken myoblast differentiation as well as iSN04. These results provide new insights into the design of shorter aptamers.

## 2. Materials and Methods

### 2.1 Oligonucleotides

PS-ODNs in which all phosphodiester bonds were phosphorothioated to enhance nuclease resistance in cell culture were synthesized and HPLC-purified (GeneDesign, Osaka, Japan) [7,16]. Unmodified oligonucleotides without phosphorothioate for nuclear magnetic resonance (NMR) and native polyacrylamide gel electrophoresis (PAGE) were synthesized and HPLC-purified (Hokkaido System Science, Sapporo, Japan). The oligonucleotide sequences are listed in Table 1. iMyo01-CS is a complementary strand DNA to iMyo01. NC-DNA is a negative control DNA that does not form a G-quadruplex, and NC-RNA-CS is a complementary strand RNA to NC-DNA.

**Table 1.**
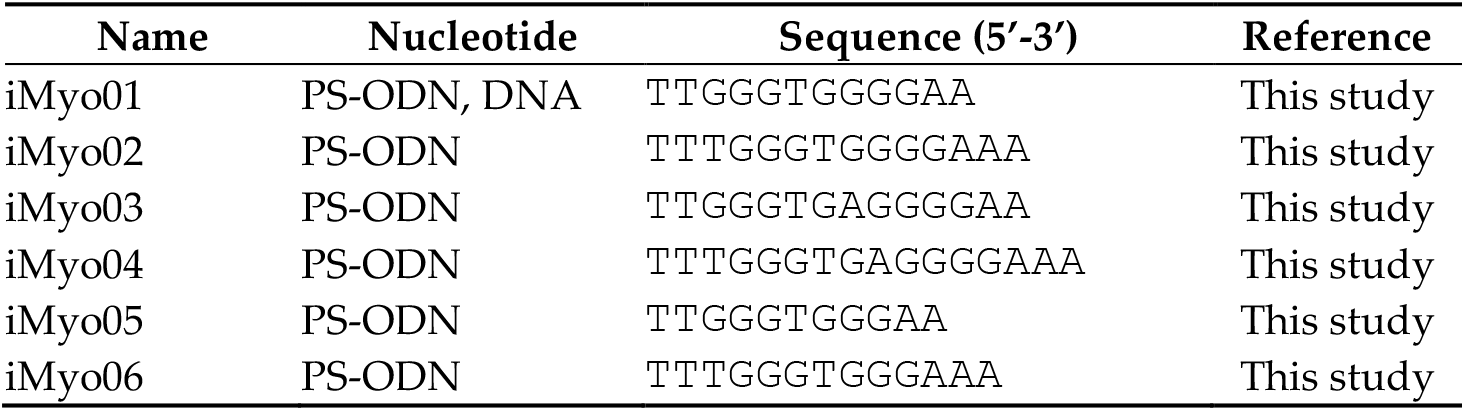
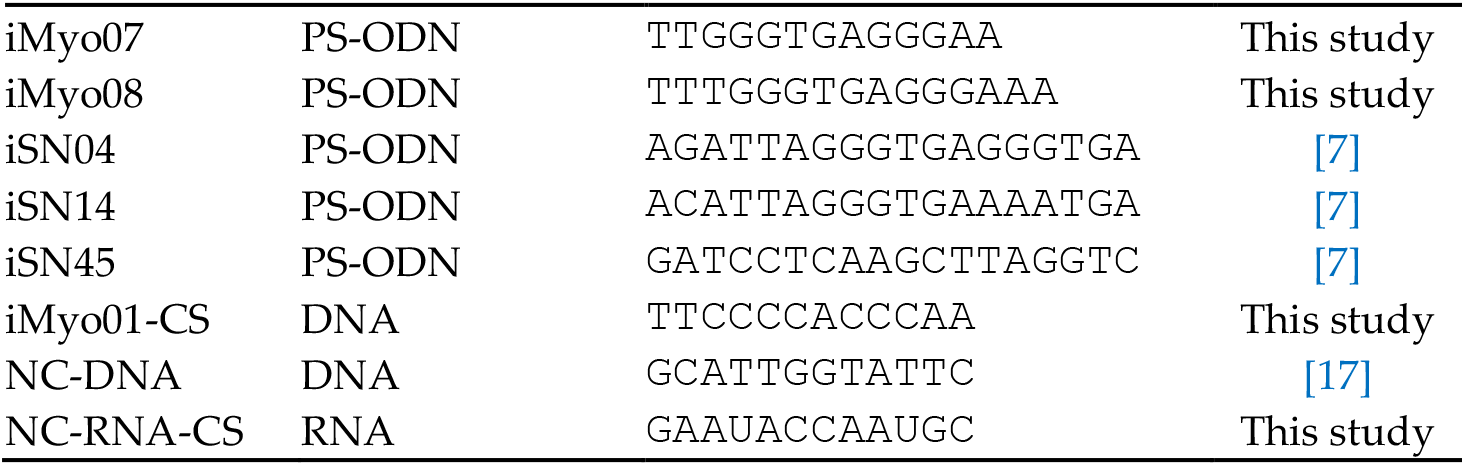
Oligonucleotide sequences.

### 2.2 Myoblast Culture

All myoblasts were cultured on the dishes/plates coated with collagen type I-C (Cell-matrix; Nitta Gelatin, Osaka, Japan) at 37°C under 5% CO_2_ throughout the experiments.

The commercially available human myoblast (hMB) stock isolated from a healthy 35-year-old female (CC-2580, lot 0000483427; Lonza, Walkersville, MD, USA) was used [7,10]. hMBs were maintained in Skeletal Muscle Growth Media-2 (CC-3245; Lonza) as a growth medium (GM) for hMB (hGM). For the screening system, 6.0×10^3^ hMBs/100 μl hGM/well were seeded on 96-well plates. For high-resolution imaging, 1.0×10^5^ hMBs were seeded on 30-mm dishes. For quantitative real-time RT-PCR (qPCR), 2.5×10^5^ hMBs were seeded on 60-mm dishes. The next day, hGM was replaced with differentiation medium (DM) for hMB (hDM) consisting of DMEM (Nacalai, Osaka, Japan), 2% horse serum (HyClone; Cytiva, Marlborough, MA, USA), and a mixture of 100 units/ml penicillin and 100 μg/ml streptomycin (P/S) (Nacalai), and PS-ODNs were treated at a final concentration of 30 μM.

Murine myoblasts (mMBs) were isolated from skeletal muscle of 4-week-old C57BL/6J mice (Clea Japan, Tokyo, Japan) as previously described [18]. Primary-cultured mMBs were maintained in GM for mMB (mGM) consisting of Ham’s F10 (Thermo Fisher Scientific, Waltham, MA, USA), 20% fetal bovine serum (FBS) (HyClone; Cytiva), 2 ng/ml recombinant human basic fibroblast growth factor (rh-bFGF) (Fujifilm Wako Chemicals, Osaka, Japan), and P/S. For the screening system, 1.0×10^4^ mMBs/100 μl mGM/well were seeded on 96-well plates. The next day, the medium was replaced with the fresh mGM containing 10 μM PS-ODNs.

Chicken myoblasts (chMBs) were isolated from the leg muscle of E10 embryos of broiler UK Chunkey chicken (National Federation of Agricultural Cooperative Associations, Tokyo, Japan) as previously described [9,19-21]. chMBs were maintained in GM for chMB (chGM) consisting of RPMI1640 (Nacalai), 20% FBS, 1% non-essential amino acids (Fujifilm Wako Chemicals), 1% chicken embryo extract (US Biological, Salem, MA, USA), 2 ng/ml rh-bFGF, and P/S. For the screening system, 5.0×10^3^ chMBs/100 μl chGM/well were seeded on 96-well plates. The next day, the medium was replaced with the fresh chGM containing 10 μM PS-ODNs.

### 2.3 Immunocytochemistry

After 48 h of PS-ODN treatment, myoblasts were subjected to immunostaining for myosin heavy chain (MHC). The myoblasts were fixed with 2% paraformaldehyde, permeabilize with 0.2% Triton X-100, and immunostained with 0.5 μg/ml mouse monoclonal anti-MHC antibody (MF20; R&D Systems, Minneapolis, MN, USA). As a secondary antibody, 0.1 μg/ml of Alexa Fluor 488-conjugated donkey polyclonal anti-mouse IgG antibody (Jackson ImmunoResearch, West Grove, PA, USA) was used. Cell nuclei were stained with DAPI (Nacalai). The screening system for analyzing MHC images has been described previously [7]. Briefly, fluorescence images were acquired automatically using CellInsight NXT (Thermo Fisher Scientific) and quantified using HCS Studio: Cellomics Scan software (Thermo Fisher Scientific) as follows. For hMBs and chMBs, MHC signal intensity was defined as the total MHC signal intensity divided by the number of nuclei. For mMBs, the ratio of MHC^+^ cells was defined as the number of nuclei in the MHC^+^ cells divided by the number of nuclei. High-resolution MHC images were obtained using EVOS FL Auto microscope (AMAFD1000; Thermo Fisher Scientific). The ratio of MHC^+^ cells was defined as the number of nuclei in all MHC^+^ cells divided by the total number of nuclei, and the fusion index was defined as the number of nuclei in multinuclear MHC^+^ myotubes divided by the total number of nuclei, which were calculated using ImageJ software (National Institutes of Health, Bethesda, MD, USA).

### 2.4 qPCR

After 24 h of PS-ODN treatment, total RNA was isolated from hMBs using Nucleo-spin RNA Plus (Macherey-Nagel, Düren, Germany) and reverse transcribed using ReverTra Ace qPCR RT Master Mix (TOYOBO, Osaka, Japan). qPCR was performed using GoTaq qPCR Master Mix (Promega, Madison, WI, USA) with StepOne Real-Time PCR System (Thermo Fisher Scientific). The amount of each transcript was normalized to that of tyrosine 3-monooxygenase/tryptophan 5-monooxygenase activation protein zeta gene (*YHHAZ*). The primer sequences were described previously [7,11]. Results are expressed as fold-change.

### 2.5 Protein Precipitation and Western Blotting

Protein precipitation by PS-ODNs was performed as described previously [7]. C2C12 murine myoblast cell line (DS Pharma Biomedical, Osaka, Japan) was maintained in DMEM supplemented with 10% FBS and P/S. Soluble whole-cell lysate of C2C12 cells was prepared using lysis buffer consisting of 0.1 M Tris-HCl (pH 7.4), 75 mM NaCl, 1% Triton X-100, and a protease inhibitor cocktail (1 mM 4-(2-aminoethyl)benzenesulfonyl fluoride hydrochloride, 0.8 μM aprotinin, 15 μM E-64, 20 μM leupeptin hemisulfate monohydrate, 50 μM bestatin, and 10 μM pepstatin A) (Nacalai). The PS-ODNs conjugated with biotin at the 5’-end (GeneDesign) were immobilized on streptavidin-coated magnetic beads (Magnosphere MS300/Streptavidin; JSR Life Sciences, Sunnyvale, CA, USA) according to the manufacturer’s instructions. 100 μg lysates and 0.6 mg iSN14-beads were mixed in 1 ml lysis buffer containing 1% NP-40 (Nacalai), and then gently rotated at 4°C overnight to eliminate non-specific proteins absorbed on PS-ODN or beads. After magnetic pulldown of iSN14-beads, the supernatants were mixed with iSN04-, iMyo01-, or iMyo03-beads and rotated at 4°C overnight. Proteins precipitated by PS-ODN-beads were dissociated in lysis buffer containing 1% NP-40, 10% glycerol, and 2% sodium dodecyl sulfate (SDS) at 95°C for 5 min. The samples were subjected to 8% SDS-PAGE followed by Western blotting using iBlot 2 Dry Blotting System (Thermo Fisher Scientific). 1.0 μg/ml rabbit polyclonal anti-nucleolin antibody (ab22758; Abcam, Cambridge, UK) and 0.1 μg/ml horseradish peroxidase (HRP)-conjugated goat anti-rabbit IgG antibody (Jackson ImmunoResearch) were used as primary and secondary antibodies, respectively. HRP activity was detected using ECL Prime reagents (GE Healthcare, Chicago, IL, USA) and captured using ImageQuant LAS 500 (GE Healthcare).

### 2.6 Agarose Gel Electrophoresis

0.8 nmol PS-ODNs and 0.8 nmol berberine hydrochloride (Nacalai) were mixed in 16 μl Ham’s F10 containing 144.1 mM Na^+^, 0.6 mM Mg^2+^, 5 mM K^+^, 0.3 mM Ca^2+^, 1 μM Fe^2+^, 1 μM Cu^2+^, and 0.1 μM Zn^2+^ cations. The mixtures were placed at 4°C overnight and then subjected to electrophoresis using TAE-buffered 3% agarose gel with 0.5 μg/ml ethidium bromide (EtBr). For monochromatic images, the gels were illuminated with 365 nm ultraviolet (UV) and the images were captured using ImageQuant LAS 500 with a 560 nm emission bandpass filter (GE Healthcare). For colored images, the gels were illuminated by 302 nm UV and the images were taken by a digital still camera without any filters to detect the 530 nm yellow emission from berberine and the 620 nm red emission from EtBr [7].

### 2.7 NMR

iMyo01 (DNA) was annealed by heating at 95°C for 5 min followed by snap-cooling on ice. Annealed iMyo01 (final concentration of 0.7 mM) was dissolved in 20 mM sodium phosphate (pH 6.5) using a centrifuge concentrator (Vivaspin 3000 MWCO; Sartorius, Göttingen, Germany). For the K^+^ condition, concentrated KCl was added to the sample to a final concentration of 50 mM. NMR spectra were measured using AVANCE600 spectrometer (Bruker BioSpin, Billerica, MA, USA) at a probe temperature of 10°C. 1D imino proton spectra and 2D nuclear Overhauser effect spectroscopy (NOESY) spectra (mixing time of 150 msec) were recorded using the jump-and-return or 3-9-19 pulse schemes for water suppression.

### 2.8 Native PAGE

The conformation of iMyo01 (DNA), iMyo01-CS, NC-DNA, and NC-RNA-CS was analyzed by 20% native PAGE with or without 5 mM KCl. Concentrated KCl solution was added to the gel and running TBE buffer. Gels were stained with SYBR Gold (Thermo Fisher Scientific).

### 2.9 Multicanonical Molecular Dynamics (TTP-McMD) Simulation

The two molecules of iMyo01 (DNA) in explicit solvent were simulated by trivial-trajectory parallelization (TTP-) McMD method [7,16]. The simulation system (Figure S1) contained two iMyo01 molecules, 10,252 water molecules, 23 K^+^ ions, and one Cl^−^ ion to be neutralized electrically. The initial structure of one iMyo01 molecule was built as a DNA helix model by NAB in AmberTools [22]. The other iMyo01 was copied from the first and was separated by 10 Å. The force field of amber OL15 [23] was applied for iMyo01. TIP3P model [24] was used for water. The periodic boundary condition was used with a truncated octahedron box. The long-range interaction was calculated by particle-meshed Ewald with 10 Å cut off.

The TTP-McMD [25] was conducted to sample the equilibrated conformations at 310 K. Before TTP-McMD, the volume of system was equilibrated by MD simulation under constant NPT condition of 1 bar. The energy range of the multicanonical ensemble covered from 270 K to 600 K. 100 trajectories were used, and 14 iterative runs were done to achieve a flat distribution along the energy range. The production run was conducted for 100 ns in each trajectory (total 10 μs). The snapshots were saved every 200 ps. The total of 50,000 snapshots were sampled, and the 988 conformations at 310 K were obtained as the 310 K canonical ensemble by reweighting method.

The representative iMyo01 structures were taken from the centroid of structural clustering using The AmberTools22 [26], which was performed for the iMyo01 molecules with the metrics of the root-mean square deviation of heavy atoms within iMyo01 in the dimer state. Hydrogen bond analysis was performed using VMD tools [27]. The structure images were generated by UCSF Chimera [28].

### 2.10 Statistical Analysis

The results were presented as mean ± standard error. Statistical comparisons were performed using multiple comparison test with Dunnett’s test following one-way analysis of variance. Statistical significance was set at *p* < 0.05.

## 3. Results

### 3.1 Designing and Screening of iMyo-ODNs

The established 18-base myoDN, iSN04, has a tandem telomeric repeat (TTA GGG TGA GGG) as the core sequence for its myogenetic activity. In particular, the G-stack consisting of the 13-15th GGG is an indispensable structure for iSN04 activity [7]. We designed eight non-telomeric PS-ODNs (iMyo-ODNs; iMyo01–iMyo08) with two G groups (GGG or GGGG) separated by a mini-linker (T or TGA), thymines at the 5’ end (TT or TTT), and adenines at the 3’ end (AA or AAA) (Table 1). It has been speculated that iMyo-ODNs form a condensed conformation through interaction between 5’-terminal thymines and 3’-terminal adenines and have G-stacks within the structure, similar to that of iSN04.

To investigate the myogenetic activities of iMyo-ODNs, undifferentiated myoblasts were treated with PS-ODNs for 48 h and immunostained for MHC, a terminal differentiation marker of skeletal muscle. The signal intensities of MHC or the ratio of MHC^+^ cells were automatically quantified in an unbiased manner using the screening system (Figure 1). iSN04 was used as a positive control, and iSN14 and iSN45 served as negative controls. In hMBs, iMyo01 and iMyo03 significantly increased MHC signal intensities to the same extent as iSN04. Other iMyo-ODNs and negative controls did not induce myogenesis (Figure 1A). In mMBs, iMyo03 and iMyo04 markedly increased the ratio of MHC^+^ cells (Figure 1B). In chMBs, iMyo01–iMyo04 exhibited significant abilities to promote myogenic differentiation (Figure 1C). In all myoblasts, iMyo05–iMyo08 did not induce myogenesis, indicating that the latter GGGG, not GGG, is essential for iMyo-ODNs to function as myoDNs.

**Figure 1.**
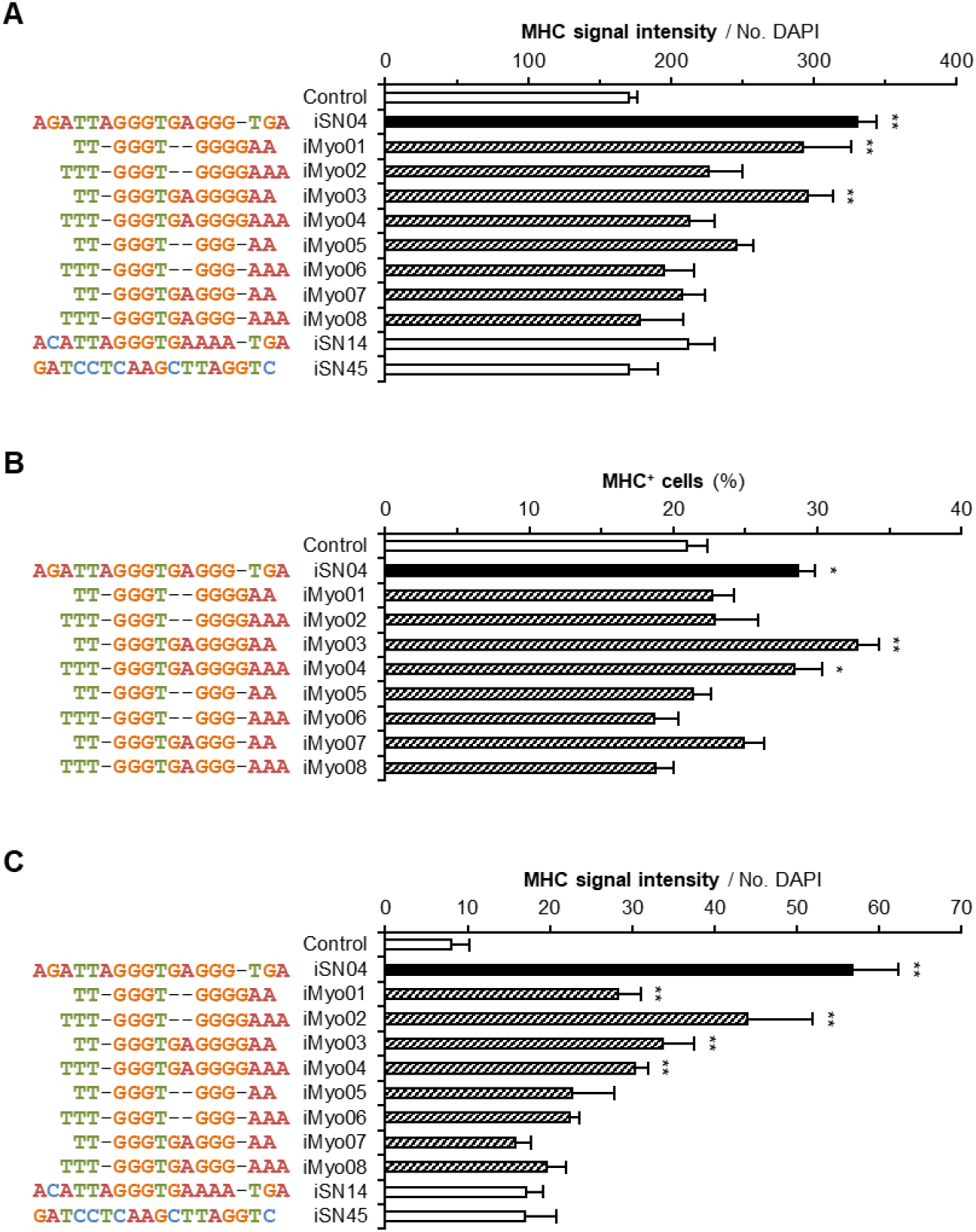
Myogenetic activities of iMyo-ODNs. **(A)** MHC signal intensities of hMBs treated with 30 μM PS-ODNs for 48 h. **(B)** The ratio of MHC^+^ cells within mMBs treated with 10 μM PS-ODNs for 48 h. **(C)** MHC signal intensities of chMBs treated with 10 μM PS-ODNs for 48 h. ^*^ *p* < 0.05, ^**^ *p* < 0.01 vs. control (Dunnett’s test). *n* = 3.

The effects of iMyo01 and iMyo03 on hMBs were further investigated by high-resolution imaging and qPCR (Figure 2). Immunostaining revealed that both iSN04 and iMyo01 significantly induced the differentiation of hMBs into MHC^+^ myocytes and multinuclear myotubes. iMyo03 also markedly accelerated myotube formation (Figure 2A). The mRNA levels of MyoD (*MYOD1*), a master regulator of myogenesis, were not altered by any of the PS-ODNs, but those of myogenin (*MYOG*), a myogenic transcription factor, were significantly increased by iMyo03 and were similarly induced by iSN04 and iMyo01.

**Figure 2.**
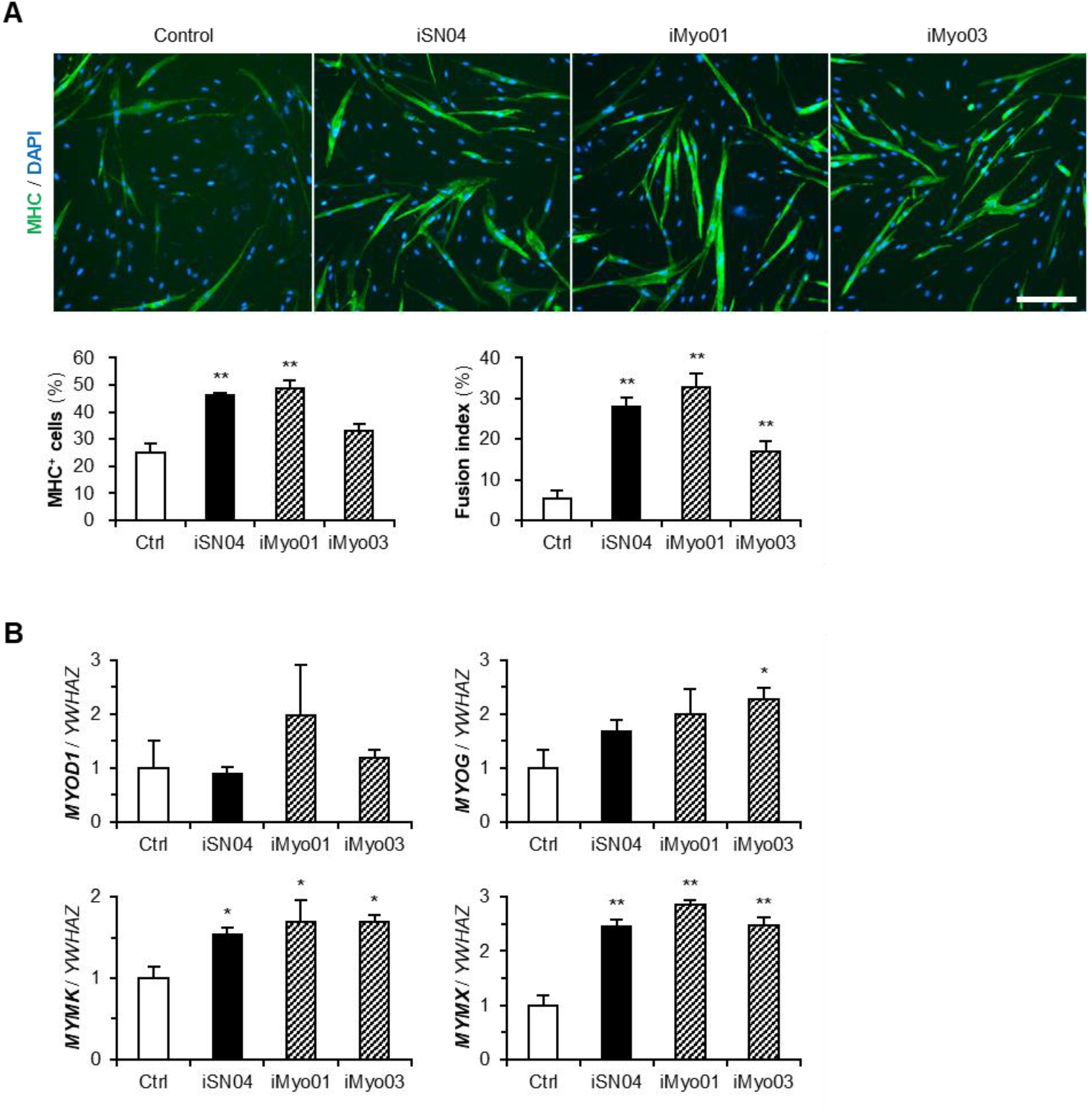
Effects of iMyo01 and iMyo03 on hMBs. **(A)** Representative immunofluorescence images of hMBs treated with 30 μM PS-ODNs in DM for 48 h. Scale bar, 200 μm. The ratio of MHC^+^ cells and multinuclear myotubes was quantified. ^**^ *p* < 0.01 vs. control (Dunnett’s test). *n* = 6-8. **(B)** qPCR results of myogenic gene expression in hMBs treated with 30 μM PS-ODNs in DM for 24 h. The mean values of control hMB were set to 1.0. ^*^ *p* < 0.05, ^**^ *p* < 0.01 vs. control (Dunnett’s test). *n* = 3-4.

Transcriptions of myomaker (*MYMK*) and myomixer (*MYMX*), which are myogenin-regulated muscle-specific membrane proteins for myotube formation, were markedly upregulated by iSN04, iMyo01, and iMyo03 (Figure 2B). These results demonstrate that iMyo01 and iMyo03 as well as iSN04 promote the myogenic differentiation of hMBs.

### 3.2 iMyo01 and iMyo03 Bind to Nucleolin But Not to Berberine

Since iSN04 physically interacts with nucleolin to exert its myogenetic activity [7], iMyo01 and iMyo03 were tested for their ability to bind to nucleolin by precipitation assay. Biotin-conjugated PS-ODNs were immobilized on streptavidin-beads. The soluble lysate of C2C12 murine myoblast cell line was pre-pulled-down with iSN14-beads to eliminate the absorption of non-specific proteins. After removing off-targets, the lysate was precipitated with iSN04-, iMyo01-, or iMyo03-beads. The precipitates were subjected to Western blotting to detect nucleolin. As shown in Figure 3A, nucleolin was precipitated by iMyo01 and iMyo03 as well as by iSN04.

**Figure 3.**
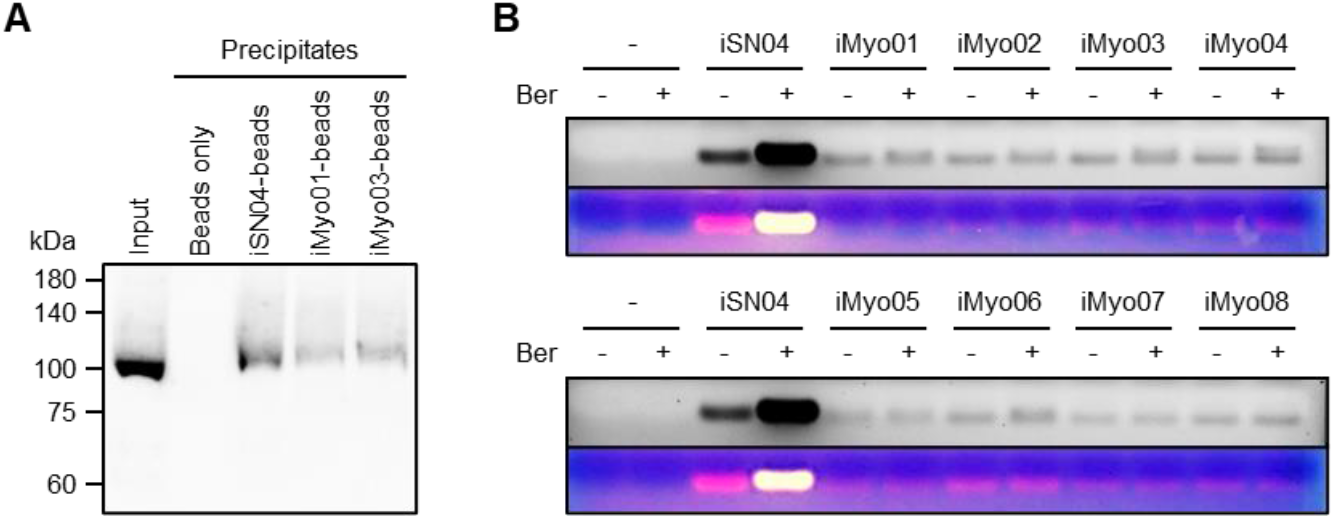
iMyo01 and iMyo03 bind to nucleolin but not to berberine. **(A)** Representative Western blotting image of nucleolin in the soluble whole-cell lysates of C2C12 cells precipitated by iSN04, iMyo01, or iMyo03. **(B)** Representative images of agarose gel electrophoresis of iSN04 and iMyo-ODNs mixed with berberine in Ham’s F10. Ber, berberine.

iSN04 also interacts with berberine via its G-stack (GGG at the 13-15th base) in the presence of Ca^2+^ or Mg^2+^ [7]. Berberine is an isoquinoline that binds to the G-quartet and stabilizes the G-quadruplex structure derived from telomeric sequences [29]. The iSN04-berberine complex shows higher myogenetic activity compared to iSN04 alone because berberine stabilizes the core G-stack within iSN04 [7]. To determine whether iMyo-ODNs form a complex with berberine, PS-ODNs and berberine were mixed in Ham’s F10 and subjected to agarose gel electrophoresis. As shown in Figure 3B, the yellow emission from berberine was detected at the same position as the red emission from iSN04 stained with EtBr, indicating the iSN04-berberine complex. However, all iMyo-ODNs, even the myogenetic iMyo01 and iMyo03, did not interact with berberine. This result suggests that the G-repeats of iMyo-ODNs form different structure than the G-stack within iSN04. Since the myogenetic activities of iMyo01 and iMyo03 undoubtedly depend on their ability to bind to nucleolin, their conformations need to be determined directly.

### 3.3. iMyo01 Forms a G-quadruplex Structure

The imino proton NMR spectra of iMyo01 (DNA) were measured to characterize its structure (Figure 4A). In the absence of KCl, seven sharp and four broad signals were observed. After titration with 50 mM KCl, the NMR spectrum changed significantly, but seven sharp and four broad signals were observed, similar to those in the absence of KCl. The imino proton resonances between 10.5 and 12.5 ppm are characteristic for G-quadruplex formation [30], consistent with that iMyo01 having no cytosine and therefore no G:C base pair. Since the G-quadruplex structure is stabilized by K^+^ [31], it is considered that the KCl-induced shift of the imino proton spectrum of iMyo01 represents the stabilization of its G-quadruplex by K^+^ binding. To confirm G-quadruplex structure, NOESY spectrum of iMyo01 was measured (Figure 4B). The strong NOE signals between imino protons and NOE signals between H8 and imino protons strongly suggested that iMyo01 forms the G-quadruplex (Figures 4B and 4C).

**Figure 4.**
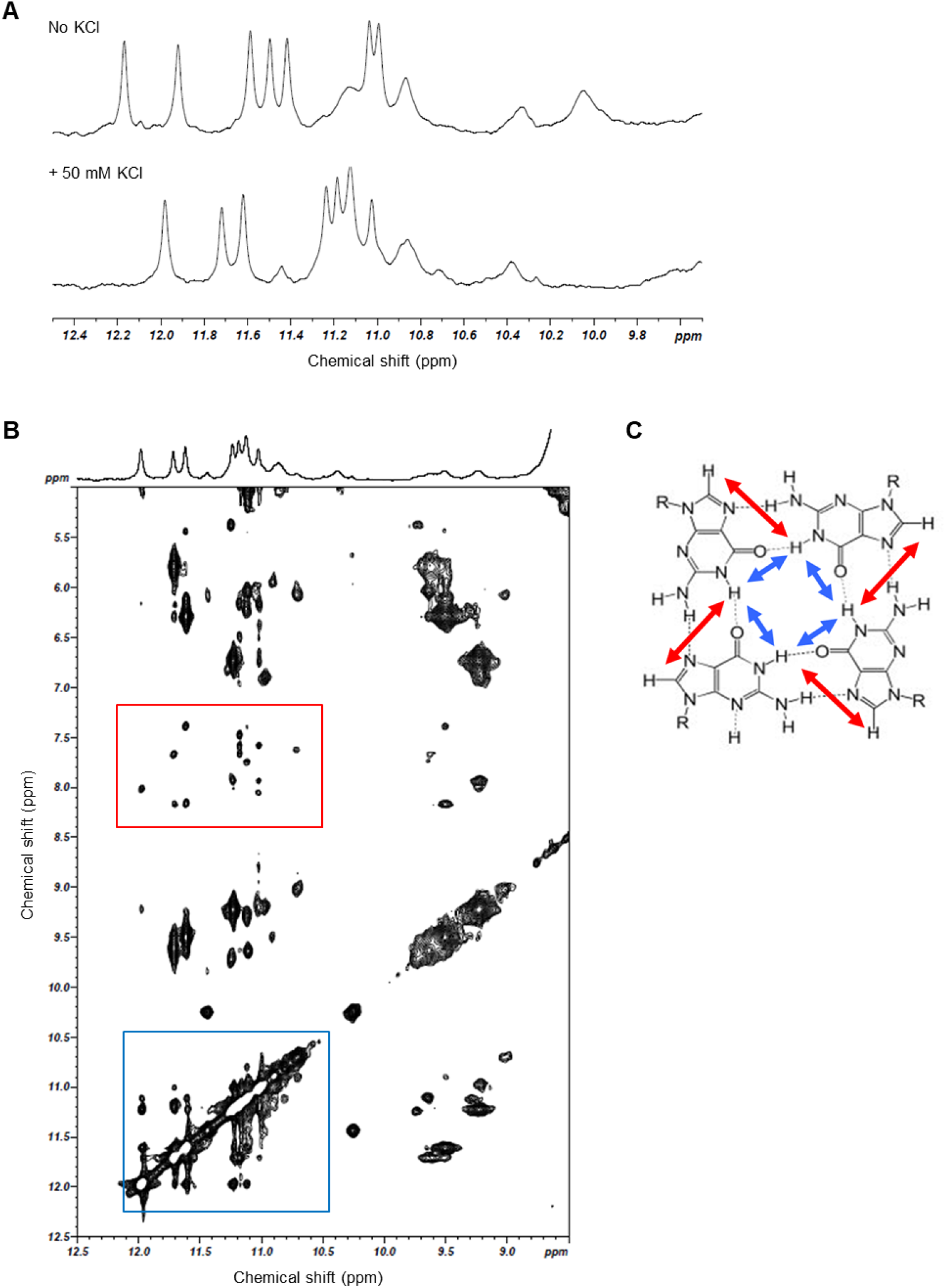
NMR analysis of iMyo01. **(A)** Comparison of 1D imino proton spectra of iMyo01 (DNA) in the absence of KCl and in the presence of 50 mM KCl. **(B)** 2D NOESY spectrum (mixing time 150 ms). The NOE signals between imino protons (blue box) and the NOE signals between imino protons and H8 (red box). **(C)** Schematic representation of G-quartet and NOEs between imino protons (blue arrows) and between imino proton and H8 (red arrows).

### 3.4. iMyo01 Forms a Homodimer in the Presence of K^+^

iMyo01, which has only two G-repeats, cannot form a G-quadruplex as a monomer. Then, the conformation of iMyo01 (DNA) was analyzed by native PAGE. As shown in Figure 5, iMyo01-CS (lane 6) was not stained with SYBR Gold due to its low number of purines (only three adenines), but other nucleotides were detected. iMyo01 (lane 4) was observed in monomer mobility in the absence of KCl, but was detected in dimer mobility in the presence of KCl, which is confirmed by the duplex of iMyo01 and iMyo01-CS (lane 5). In contrast, no dimer mobility was observed for NC-DNA (lane 1), which never forms a G-quadruplex, even in the presence of KCl. The K^+^-dependent dimer formation strongly suggests that iMyo01 forms a G-quadruplex structure in the presence of K^+^, in agreement with the results of the NMR.

**Figure 5.**
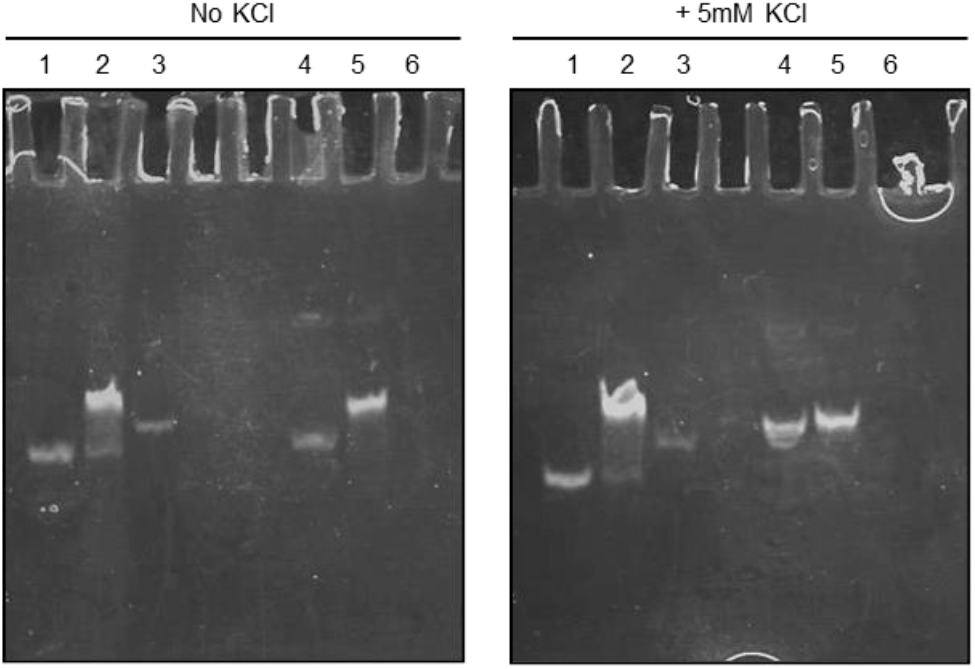
Native PAGE of iMyo01. Lane 1, NC-DNA; lane 2, duplex of NC-DNA and NC-RNA-CS; lane 3, NC-RNA-CS; lane 4, iMyo01 (DNA); lane 5 duplex of iMyo01 (DNA) and iMyo01-CS; lane 6, iMyo01-CS.

Dimer formation of iMyo01 (DNA) was computationally simulated by McMD with the all-atom model. Using the simulated conformational ensemble, the distances between the centers of mass of the two iMyo01 molecules (*d*_com_) were calculated. Their probability distributions showed that the dimer state (*d*_com_ < 25 Å) is more stable than the separated state (*d*_com_ ≥ 25 Å) (Figure S2). The conformation at *d*_com_ ≈ 10 Å had the highest probability, which was the most stable. Structural clustering of the dimer state represented the centroid structures of the top three clusters occupying 86.5% of all structures. Their *d*_com_ values were 6.0 (Figure 6A), 11.4 (Figure 6B), and 18.6 Å (Figure 6C), suggesting that the conformation in the second cluster can be the stable dimer. The analysis of the intra/inter-molecular hydrogen bonds between the residues of iMyo01 showed that the guanines form hydrogen bonds. The distinct hydrogen bonds among the G-repeats in the second cluster may correspond to the G-quadruplex detected by NMR.

**Figure 6.**
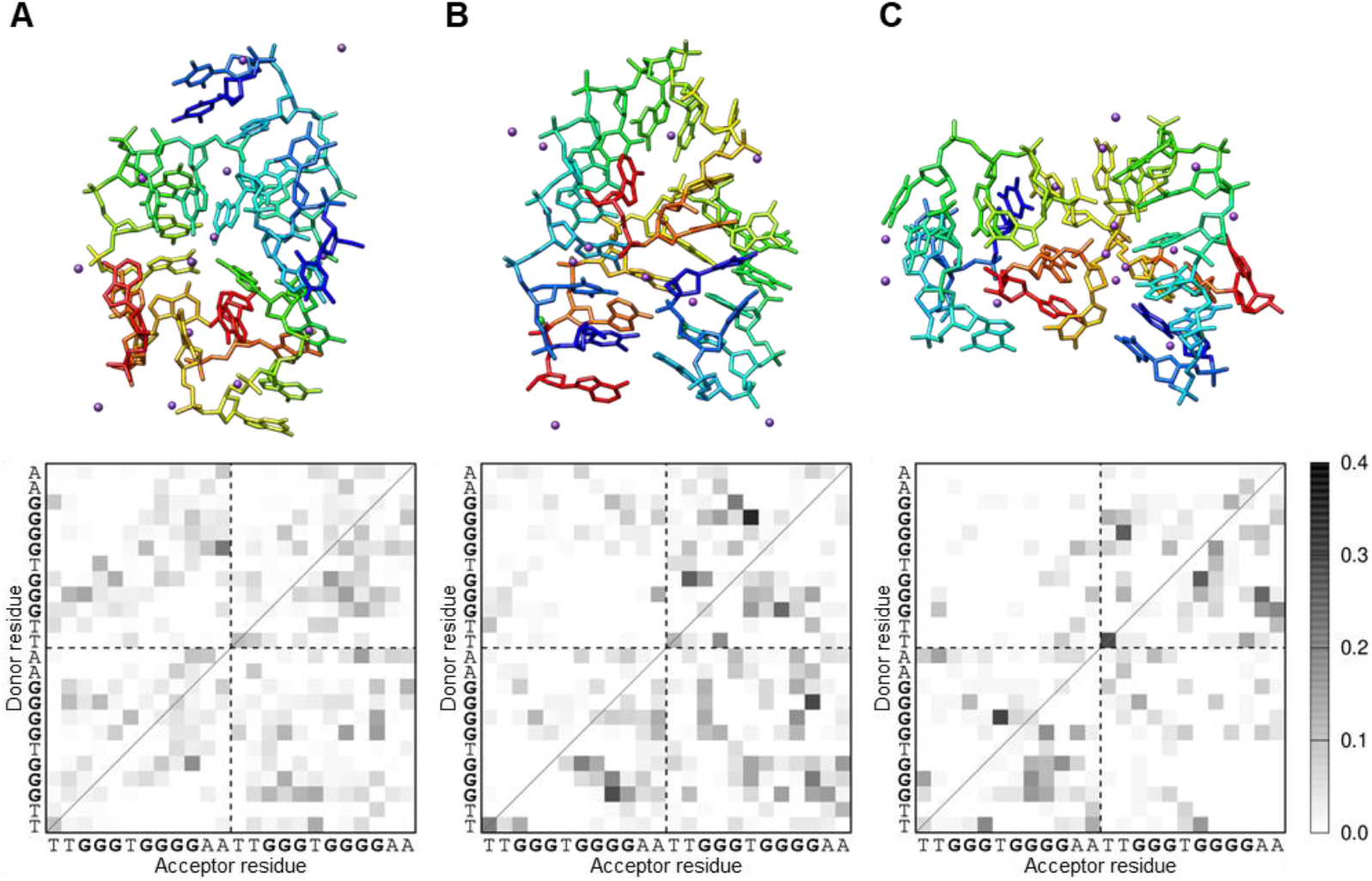
The representative iMyo01 structures in the dimer state from McMD simulation. (**A**) The first cluster (42.1%; *d*_com_, 6.0 Å). (**B**) The second cluster (27.0%; *d*_com_, 11.4 Å). (**C**) The third cluster (17.5%; *d*_com_, 18.6 Å). The clusters are shown as a rainbow-colored stick model from the 5’ to the 3’ end. The K^+^ ions are shown as purple spheres. The hydrogen bond patterns in each cluster are plotted as a monochrome map. The gray scale indicates the probability of hydrogen bond formation between residues. The two iMyo01 sequences are displayed sequentially on the axes.

The hydrogen-bond patten within iMyo01 showed that the G3-G5 can interact with the G7-G10. The intra hydrogen-bond pattern can be formed in the separated monomer state (Figure S3). The monomer state had the stacking tandem guanines. In the second cluster of the dimer state, the intramolecular hydrogen bonds were maintained as well as in the monomer state. Furthermore, the K^+^ ions were found at the interface of iMyo01. The simulation results also indicated that the K^+^ was important in mediating the two iMyo01 molecules to form the dimer, which might reduce electrostatic repulsion between them. However, the G-stack in the dimer was slightly perturbed so that the simulated conformations did not have a fully coplanar arrangement of four guanines, suggesting that a more accurate force field of nucleic acids and solvents would be required to reproduce the complete G-tetrad.

## 4. Discussion

The present study successfully developed the novel shorter myoDNs that promote myogenic differentiation. iMyo03 induced differentiation of hMBs, mMBs, and chMBs as well as the established myoDN, iSN04. iMyo01 activated hMBs and chMBs, whereas iMyo02 and iMyo04 affected only chMBs. Such species-specificity of iMyo-ODNs may be due to the differences of nucleolin structures. Although myoDNs are considered to interact with four RNA-binding domains (RBD1-RBD4) of nucleolin [9], the amino acid sequences of the nucleolin RBDs are not completely conserved among animals (Table 2). The previous study have also reported that G-quadruplex anti-thrombin DNA aptamers (15-TBA, 31-TBA, and RA-36) individually exhibited different activities against various mammalian thrombins [32]. Clinically, aptamer cross-reactivity with the target ortholog in a patient can lead to side effects [33]. Our results suggest that modification of the existing aptamer can alter not only the affinity but also the cross-reactivity to the target molecule, which needs to be considered in aptamer design.

**Table 2.** Amino acid sequence identities of nucleolin RBDs.

| RBD | Human vs.<br>mouse | Human vs.<br>chicken | Mouse vs.<br>chicken |
| --- | --- | --- | --- |
| RBD1 | 85.9% | 56.4% | 56.4% |
| RBD2 | 83.8% | 69.3% | 64.5% |
| RBD3 | 89.3% | 71.1% | 67.1% |
| RBD4 | 93.7% | 88.2% | 89.5% |

The 12-base iMyo01 is the shortest myoDN found and is two-thirds the length of the established 18-base iSN04. Both iMyo01 and iSN04 act as nucleolin-binding aptamers, their lengths are much shorter than the average of known aptamers (51 bases) [34]. This is an advantage of our myoDNs as nucleic acid drugs. Because ODNs are chemically synthesized by sequential coupling of phosphoramidite monomers, longer ODNs have several issues related to purity, cost, and scalability for industrial mass production [35]. As biomolecules, longer ODNs have a greater chance of being phagocytized and cross-recognized [15], compromising drug efficacy and safety. Therefore, shortening aptamers while maintaining their activities is an important and beneficial challenge in the development of aptamers as nucleic acid drugs. In addition, myoDNs must be taken up into cells because their target protein, nucleolin, is located in the nuclei. However, iSN04 is incorporated into the cytoplasm without a carrier [7], such gymnosis generally occurs with ODN treatments [14]. Since myoDNs need to cross plasma, endosomal, and nuclear membranes to reach nucleolin in the nucleus, the reduction in molecular weight will also contribute to their intracellular uptake.

On the other hand, the myogenetic activity of iMyo01 on hMBs was almost cell-physiologically equivalent to that of iSN04. The working doses of iSN04 and iMyo01 need to be carefully evaluated in further studies because iMyo01 acts as a homodimer. Although the nucleolin-binding ability of iMyo01 appeared to be lower than that of iSN04 in the precipitation assay, this is probably due to the inability of iMyo01 immobilized on the beads to efficiently form homodimers.

Both experimental and computational results indicated that iMyo01 forms a G-quadruplex structure by dimerization in the presence of K^+^. Unmodified DNA molecules were used for these cell-free structural analyses due to the absence of nucleases. The nuclease-resistant PS-ODNs were used for cellular experiments, but they are mixtures of enantiomers of phosphorothioates and are not suitable for conformational analysis. Although the formation and stabilization of tertiary structures of nucleic acids are possibly affected by nucleotide modifications, UV thermodynamic analysis has shown that phosphorothioates do not have a significant effect on the stability of the G-quadruplex structure [36]. Therefore, the conformation of unmodified iMyo01 confirmed by NMR and molecular simulation would essentially match that of PS-iMyo01. It is still difficult to selectively synthesize the designated enantiomer of PS-ODNs, and also hard to determine its structure experimentally. In silico calculations of configured enantiomers can be a powerful approach to overcome this problem and improve the productivity of aptamer design. The identification of iMyo01 in this study demonstrated that combined research technologies are useful for functional shortening of aptamers. It will hopefully contribute to the development of nucleic acid drugs.

## 5. Conclusions

This study developed the 12-base short dimeric myoDN, iMyo01, which serves as an anti-nucleolin aptamer to promote myoblast differentiation, as well as the well-known 18-base iSN04. iMyo01 dimerizes in the presence of K^+^ to form a G-quadruplex structure, resulting in nucleolin-binding and myogenetic capabilities. This is a successful instance of shortening the aptamer while maintaining its bioactivity, providing useful insights for the development of aptamers as nucleic acid drugs.

## 6. Patents

K.U. and T.T. are the inventor of Japanese Patent No. 7386507 covering iMyo-induced myogenic differentiation.

### Supplementary Materials

Figure S1: The simulation system of the iMyo01 (DNA) dimer; Figure S2: The probability of the center-of-mass distances between the two iMyo01 molecules; Figure S3: The representative iMyo01 structure in the separated monomer state.

### Author Contributions

Conceptualization, K.U. and T.T.; investigation, K.U., R.I., T. Sakamoto, Y.E., Y.N., S.S. and T.T.; resources, T. Shimosato. and H.K.; writing—original draft preparation, K.U., T. Sakamoto., and T.T.; funding acquisition, K.U., T. Sakamoto, T.T

### Funding

This research was funded by the Japan Society for the Promotion Science (19K05948 and 22K05554) to T.T.

### Institutional Review Board Statement

The experimental procedure for the preparation of mMBs and chMBs were performed in accordance with the Regulations for Animal Experimentation of Shinshu University, and the animal protocol was approved by the Committee for Animal Experiments of Shinshu University (No. 280083).

### Informed Consent Statement

The hMBs used in this study is commercially available. Detailed information of hMB is described in certificate analysis; https://bioscience.lonza.com/lonza_bs/CH/en/coa/search

### Data Availability Statement

The data presented in this study are available on request to the corresponding author.

### Conflicts of Interest

The authors declare no conflict of interest.

---

**Supplementary Figure S1.**
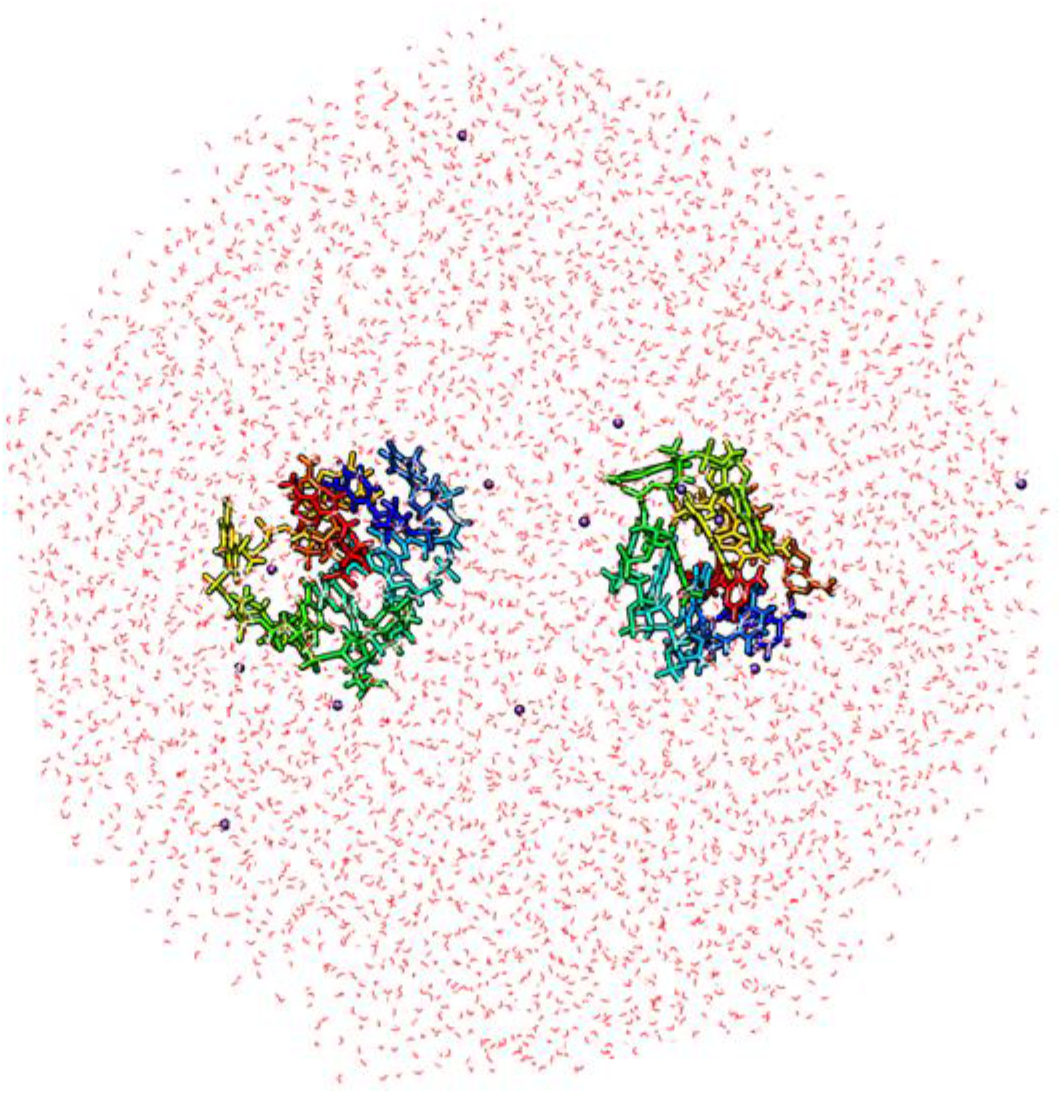
The simulation system of the iMyo01 (DNA) dimer. iMyo01 molecules are represented as a rainbow-colored stick model from the 5’ to the 3’ end. The solvent molecules are shown as red wires (oxygen atoms of water molecules), purple spheres (K^+^ ions), and a green sphere (Cl^-^ ion).

**Supplementary Figure S2.**
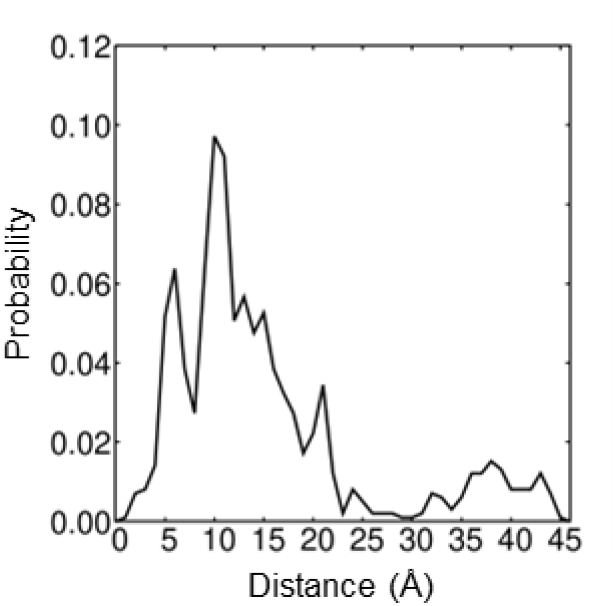
The probability of the center-of-mass distances between the two iMyo01 molecules. The McMD simulation at 310 K provided 988 conformations as a canonical ensemble. Their center-of-mass distances and probability distribution are shown. The dimer state defined below 25 Å of the distance contains 855 conformations (86.5%), and the separated monomer state contains 133 conformations (13.5%).

**Supplementary Figure S3.**
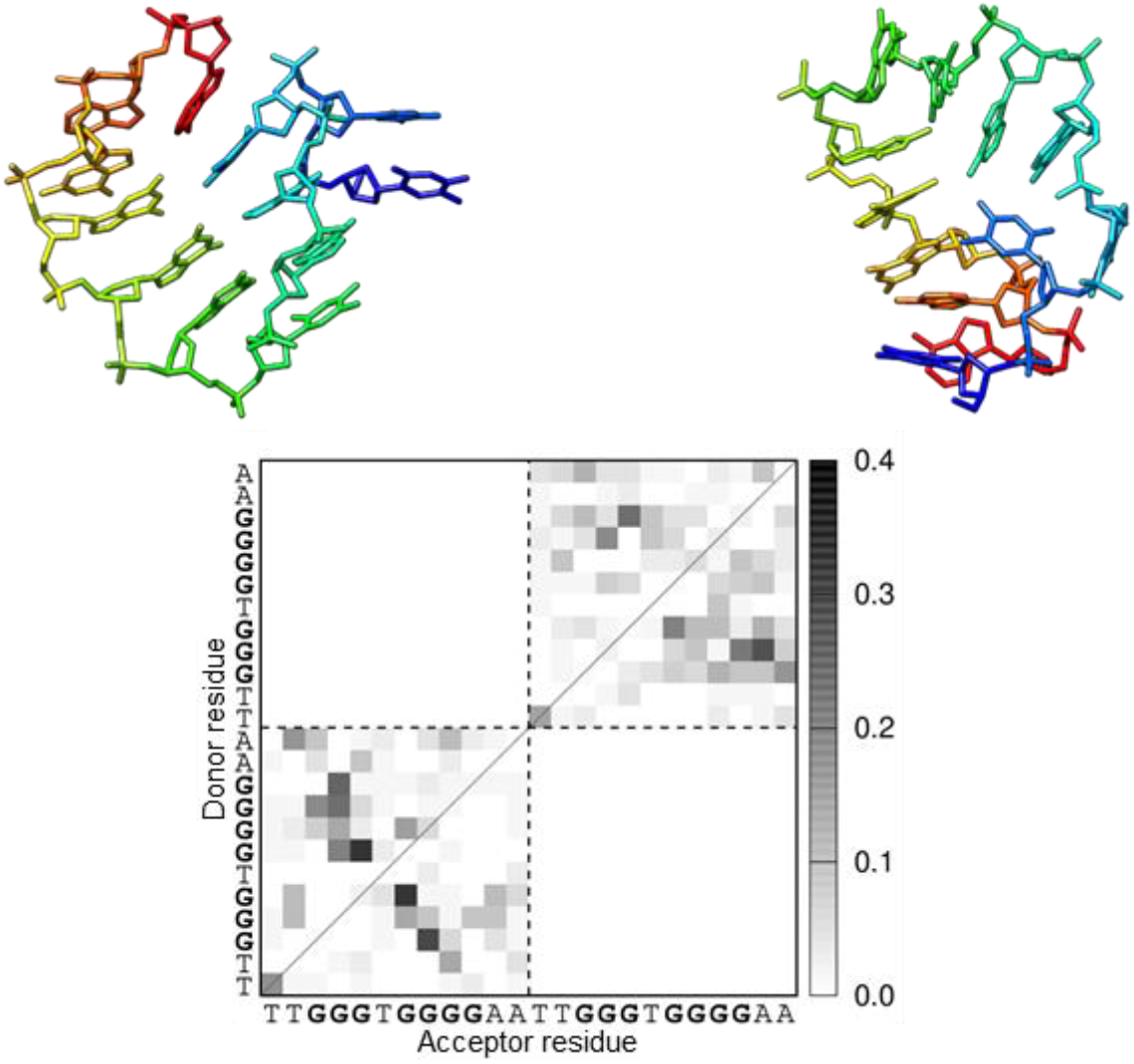
The representative iMyo01 structure in the separated monomer state. The structures (*d*_com_, 37.8 Å) are shown as a rainbow-colored stick model from the 5’ to the 3’ end. The hydrogen bond pattern is plotted as a monochrome map. The gray scale indicates the probability of hydrogen bond formation between residues. The two iMyo01 sequences are displayed sequentially on the axes.

## Notes

### Competing Interest Statement

The authors have declared no competing interest.

